# Spatial patterns of *Hyalomma marginatum*-borne pathogens in the Occitanie region (France), a focus on the intriguing dynamics of *Rickettsia aeschlimannii*

**DOI:** 10.1101/2024.03.05.583534

**Authors:** Joly-Kukla Charlotte, Bernard Célia, Bru David, Galon Clémence, Giupponi Carla, Huber Karine, Jourdan-Pineau Hélène, Malandrin Laurence, Rakotoarivony Ignace, Riggi Camille, Vial Laurence, Moutailler Sara, Pollet Thomas

## Abstract

*Hyalomma marginatum* is an invasive tick species recently established in mainland southern France. This tick is known to host a diverse range of human and animal pathogens such as *Rickettsia aeschlimannii*, *Theileria equi*, *Anaplasma phagocytophilum*, *Anaplasma marginale*, *Ehrlichia minasensis*, Crimean-Congo hemorrhagic fever and West Nile virus. While information about the dynamics of these pathogens is crucial to assess disease risk and develop effective monitoring strategies, few data on the spatial dynamics of these pathogens are currently available. We thus collected ticks in 27 sites in the Occitanie region to characterize spatial patterns of *H. marginatum*- borne pathogens. Several pathogens have been detected: *Theileria equi* (9.2%), *Theileria orientalis* (0.2%), *Anaplasma phagocytophilum* (1.6%), *Anaplasma marginale* (0.8%) and *Rickettsia aeschlimannii* (87.3%). Interestingly, we found a spatial clustered distribution for the pathogen *R. aeschlimannii* between two geographically isolated areas with infection rates and bacterial loads significantly lower in Hérault/Gard departments (infection rate 78.6% in average) compared to Aude/Pyrénées-Orientales departments (infection rate 92.3% in average). At a smaller scale, *R. aeschlimannii* infection rates varied from one site to another, ranging from 29% to 100%. Overall, such high infection rates (87.3% in average) and the effective maternal transmission of *R. aeschlimannii* might suggest a role as a tick symbiont in *H. marginatum*. Moreover, currently identified as a human pathogen, such results also question about its pathogenic status in humans given the low number of human cases. Further studies are thus needed to understand both the status and the role of *R. aeschlimannii* in *H. marginatum* ticks.

**IMPORTANCE:** Ticks are obligatory hematophagous arthropods which transmit pathogens of medical and veterinary importance. Their infections cause serious health issues in humans and considerable economic loss in domestic animals. Information about the presence of pathogens in ticks and their dynamics is crucial to assess disease risk for public and animal health. Analysing tick-borne pathogens in ticks collected in 27 sites in the regions Occitanie, our results highlight clear spatial patterns in the *Hyalomma marginatum*-borne pathogen distribution and strengthen the postulate that it is essential to develop effective monitoring strategies and consider the spatial scale to better characterize the circulation of tick-borne pathogens.

## INTRODUCTION

Zoonoses are responsible for 60% of emergent diseases (WHO, 2023). Ticks play an important role in the spread of zoonotic diseases and are considered the primary vectors of pathogens in Europe (Boulanger et al. 2019). In France, there are about 45 tick species of interest for public and veterinary health, including *Ixodes* spp., the main vector of pathogens responsible for the Lyme disease and anaplasmosis (Pérez-eid 2007).

*H. marginatum* is a tick species that has recently become established in mainland southern France (Vial et al. 2016) although it has been established for decades on the French Mediterranean island of Corsica (Delpy 1946; Morel 1959; Grech-Angelini et al. 2020). *H. marginatum* is currently endemic in several countries from the Maghreb to the Iberian Peninsula and the eastern Mediterranean basin, including Turkey and the Balkans (ECDC 2023). With the increase of temperatures due to climate change, this tick species may become established in northern latitudes via animal movements and/or bird migrations (Gray et al. 2009; Vial et al. 2016; Fernández-Ruiz and Estrada-Peña 2021). It appears that *H. marginatum* become established in regions with specific climate features such as warm temperatures in summer and low precipitation which are typical of the Mediterranean climate (Bah et al. 2022; Estrada-Peña, Martínez Avilés, and Muñoz Reoyo 2011). In mainland France, *H. marginatum* establishment has been reported in several departments including Pyrénées-Orientales, Aude, Hérault, Gard, Var, Ardèche, Drôme (Bah et al. 2022). Since an invasion process seems to happen in the south of France, its adaptation abilities and its current expansion area in France are being closely monitored.

As many other tick species, *H. marginatum* harbours complex microbial communities, collectively known as the microbiota encompassing symbionts, commensals, environmental microbes and on the other hand, pathogens affecting vertebrate hosts. Among the pathogens, *H. marginatum* can carry a large diversity of bacteria, viruses and parasites, in Eurasia and Africa (Bonnet et al. 2023). In Afrotropical regions, Mediterranean basin and more particularly in France, *H. marginatum* is considered as the main candidate for the transmission of the deadly Crimean-Congo hemorrhagic fever virus (CCHFV) (Bernard et al. 2022; Perveen and Khan 2022). Besides, CCHFV was detected for the first time in France in ticks from this study and in ticks collected in 2023 (Bernard et al. 2024).

The risk represented by this tick requires an exhaustive identification of pathogens it carries and the characterization of their dynamics in both space and time. Based on previous studies, *H. marginatum* distribution in the Occitanie region (NUTS-1) seems to be clustered between two geographic areas in Gard and northern Hérault departments and on the other hand Pyrénées-Orientales and south Aude departments (Bah et al. 2022). No *H. marginatum* was found so far between these two geographic clusters for unclear reasons. In this context, we hypothesized that the spatial distribution of microbial communities particularly pathogens might vary depending on the geographic cluster. It is now accepted that spatial patterns at different scales impact tick distribution, density and their associated pathobiome, due to the of environmental characteristics such as abiotic factors (temperature, landscape, vegetation) and the presence of hosts in given areas forming specific environmental niches (Pollet et al. 2020; Bonnet and Pollet 2021). The small spatial scale is important to consider as the pathogen prevalence can vary among a given biotope, probably due to specific environmental factors (Sormunen et al. 2018). At a larger spatial scale, the type of habitat can also participate to the prevalence of pathogens as it was the case for *A. phagocytophilum* and *Rickettsia* sp. of the spotted fever group infections in *Ixodes ricinus* collected in Central France from two different habitats (pasture vs woodland) (Halos et al. 2010). Another study reported differences in *Borrelia burgdorferi* prevalences between urban/suburban habitats and natural/agricultural in Slovakia (Kazimírová et al. 2023).

While a previous study identified *H. marginatum*-borne pathogens in several sites of collection in South of France between 2016 and 2019 (Bernard et al. 2024), information about the dynamics of these pathogens remain particularly scarce. We thus propose in this study a focus on the spatial distribution of *H. marginatum*-borne pathogens with a focus on its clustered distribution in Occitanie, and by taking into account the influence of others factors such as the tick sex and the engorgement status (Sperling et al. 2020; Ahantarig et al. 2013; Dergousoff and Chilton 2012; Gern, Zhu, and Aeschlimann 1990).

To this purpose, a large-scale tick collection program was performed in the Occitanie region in May 2022 and resulted in the analysis of 510 ticks found in 27 sites across four departments. We characterized the influence of spatial patterns, tick sex and engorgement status on tick-borne pathogens infection rates and loads.

## MATERIALS AND METHODS

### Study area and tick collection

A large-scale tick sampling was conducted during two weeks in May 2022. From the Stachurski and Vial (2018) study, we identified and visited 42 sites (riding schools and farms) from six French departments of the Occitanie region (**Figure 1**). Ticks were morphologically identified using a binocular loupe, sorted by sex (male/female) and engorgement status (fed, semi-engorged and unfed) and then stored at - 80°C until further use. After the identification, a total of 510 *H. marginatum* ticks were analysed. They came from four departments of the Occitanie region: Hérault, Gard, Aude and Pyrénées-Orientales whereas it was not detected in Tarn and Haute-Garonne departments (**Figure 1**). We chose a maximum of 34 ticks per site, with an average of 18.8 ticks per site. When the number of ticks was lower than 34, all ticks were analyzed. When possible, an equal proportion of males and females were selected for each site.

**Figure 1:**
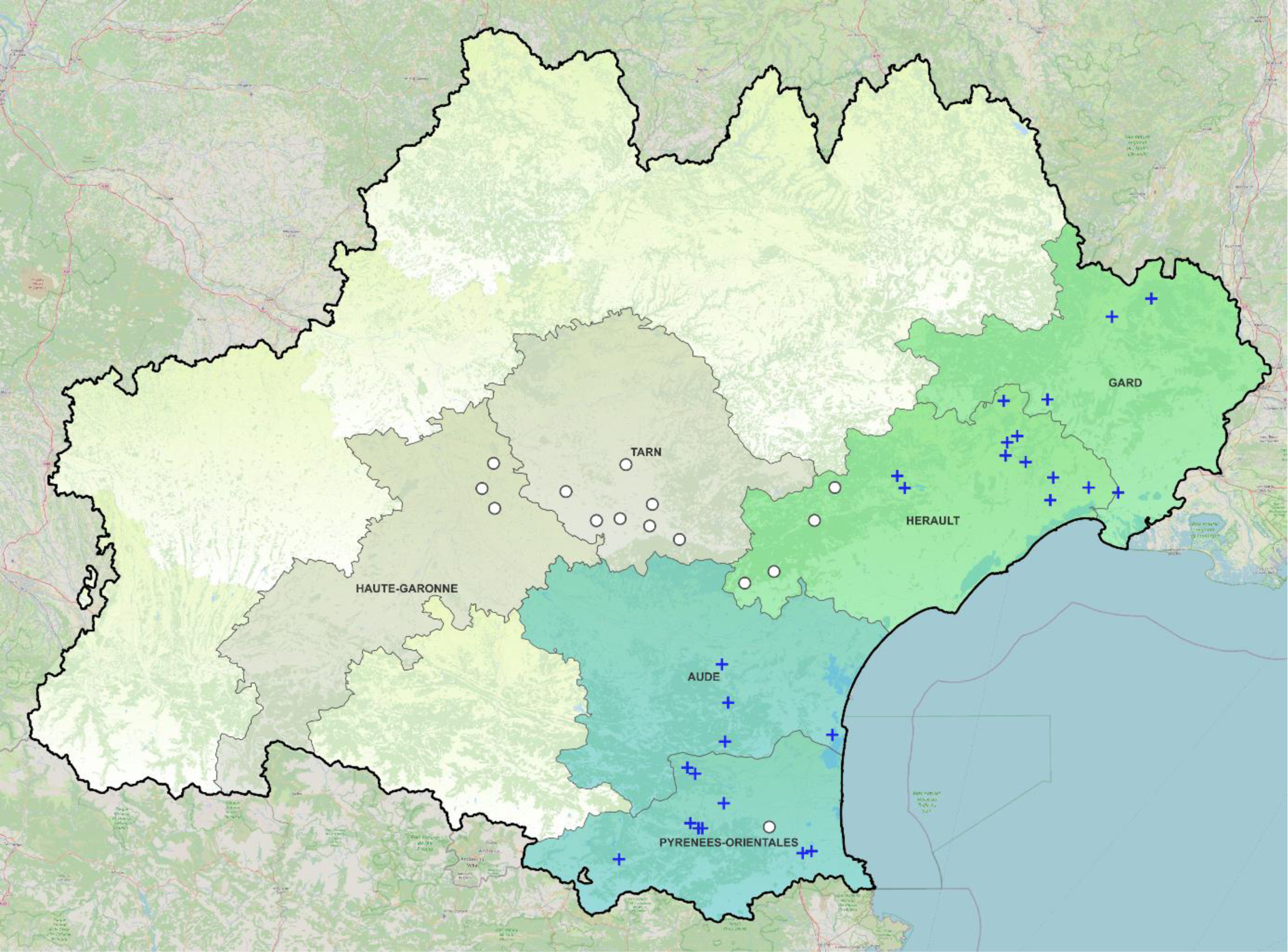
Map of the region Occitanie where the tick sampling was performed in May 2022. The Occitanie region is delimited by a black line. White circles indicate *H. marginatum* free sites and blue crosses indicate *H marginatum* positive sites where ticks were collected and selected for analyses in 2022. Blue-coloured departments represent the geographic cluster Aude/Pyrénées-Orientales and the green-coloured departments represent the geographic cluster Hérault/Gard. Grey-coloured departments were visited but no *H. marginatum* were found. Uncoloured areas correspond to other departments that were not visited for the tick collection since *H. marginatum* introduction/installation was not reported (ECDC).

### DNA and RNA extraction

Tick crushing and nucleic acids extraction were done in a BSL3 facility. Ticks were washed in bleach for 30 seconds then rinsed three times for one minute in milli Q water to limit eliminate the environmental microbes present on the tick cuticle (Binetruy et al. 2019). Ticks were then cut using a scalpel blade and crushed individually in the homogenizer Precellys®24 Dual (Bertin, France) at 5,500 rpm for 40 sec, using three steel beads (2.8 mm, OZYME, France) in 400 µL of DMEM (Dulbecco’s Modified Eagle Medium, Eurobio Scientific France) with 10% foetal calf serum. Total DNA and RNA was extracted using the NucleoMag VET extraction kit (Macherey-Nagel, Germany) as described by the manufacturer’s instructions with the IDEAL^TM^ 96 extraction robot (Innovative Diagnostics, France). Nucleic acids were eluted in 90 µL of elution buffer and stored at -20°C for DNA and -80°C for RNA until further analyses.

### Tick-borne pathogens detection in tick DNA and RNA Microfluidic PCR detection

#### Reverse transcription

Sample were retrotranscribed in cDNA using 1 µL of RNA with 1 µL of Reverse Transcription Master Mix and 3 µL of RNase-free ultrapure water provided with the kit (Standard Biotools, USA) using a thermal cycler (Eppendorf, Germany) with the following cycles: 5 min at 25°C, 30 min at 42°C and 5min at 85°C with a final hold at 10°C.

#### Targeted tick-borne pathogens

The tick-borne pathogens with a high probability to be carried by *H. marginatum* were targeted using 48 sets of primers (designs), according to data mining of the available literature (Bernard et al. 2024); Michelet et al. 2014; Gondard et al. 2018) (**Table 1**). On the 48 sets of designs, 14, 5 and 27 targeted bacteria, parasites and viruses respectively. Two designs were used for positive controls. The 48 designs were pooled to reach a 200 nM final concentration for each primer. CCHFV detection on samples was performed apart from this study (Bernard et al. 2024).

**Table 1:** Tick-borne pathogens targeted and the specific genes (designs) using the high-throughput microfluidic system (Biomark^TM^ assay).

| Genus / Family | Species | Targeted gene |
| --- | --- | --- |
| *Anaplasma* | *A. marginale* | *msp1b* |
|  | *A. phagocytophilum* | *msp2* |
|  | *Anaplasma* spp. | 16S rRNA |
| *Bartonella* | *B. henselae* | *Pap31* |
|  | *Bartonella* spp. | *ssrA* |
| *Borrelia* | *B. miyamotoi* | *glpQ* |
|  | *Borrelia* spp. | 23S rRNA |
| *Coxiella* | *Coxiella burnetii* and *Coxiella*-like endosymbionts | *idc*<br>*IS1111* |
| *Ehrlichia* | *Ehrlichia* spp. | 16S rRNA |
| *Francisella* | *F. tularensis* | *tul4* |
|  | *Francisella*-like endosymbionts | *fopA* |
| *Rickettsia* | *R. aeschlimannii* | 23s-5S ITS |
|  | *Rickettsia* spp. | *gltA* |
| *Babesia* | *B. caballi* | *Rap1*/48kDa mero antigen |
|  | *B. canis* | 18S rRNA |
| *Theileria* | *T. equi* | *ema1* |
|  | *T. annulata* | 18S rRNA |
|  | *Theileria* spp. | 18S rRNA |
| *Bunyaviridae* | Nairobi sheep disease virus | segment M |
| *Flaviviridae* | West Nile Virus | polyprotein gene |
|  | Alkhurma virus | C gene |
| *Nairoviridae* | Hazara virus | segment L |
| *Orthomyxoviridae* | Thogoto virus | segment 6 |
|  | Dhori virus | segment 2 |
| *Phenuiviridae* | Bhanja virus | *Pol*, N |

See Michelet et al. 2014; Gondard et al. 2018b; 2020; Crivei et al. 2023; Bernard et al. 2024 for more details.

#### Pre-amplification

Each sample was pre-amplified using 1.25 µL of DNA mix (1:1 volume ratio of DNA and cDNA) with 1 µL of the PreAmp Master mix (Standard Biotools, USA), 1.5 µL of ultra-pure water and 1.25 µL of the pooled designs. PCRs were performed using a thermal cycler with the following cycles: 2 min at 95°C, 14 cycles at 95°C for 15 sec and 60°C for 4 min and finally a 4-min hold at 4°C. A negative control was used for each plate with ultra-pure water. Amplicons were diluted 1:10 with ultra-pure water and stored at -20°C until further use.

#### BioMark^TM^ assay

The BioMark^TM^ real-time PCR system (Standards Biotools, USA) was used for high-throughput microfluidic real-time PCR amplification using the 48.48 dynamic array (Standard Biotools, USA). The chips dispense 48 PCR mixes and 48 samples into individual wells, after which on-chip microfluidics assemble PCR reactions in individual chambers prior to thermal cycling resulting in 2,304 individual reactions. In one single experiment, 47 ticks and one negative control are being tested. For more details, please see (Michelet et al. 2014; Gondard et al. 2018a).

#### Validation of the results by PCR and sequencing

Conventional PCRs or qPCR were then performed for tick-borne pathogens positive-samples using different sets of primers than those used in the BioMark^TM^ assay to confirm the presence of pathogenic DNA (**Table S1**). Amplicons were sequenced by Azenta Life Sciences (Germany) using Sanger-EZ sequencing and assembled using the Geneious software (Biomatters, New-Zealand). An online BLAST (National Center for Biotechnology Information) was done to compare results with published sequences in GenBank sequence databases.

### Detection and quantification of *T. equi* and *R. aeschlimannii* by duplex Real-Time Fluorescence Quantitative PCR

Tick samples were also screened for the detection and quantification of *T. equi* and *R. aeschlimannii* using a second detection method by qPCR with primers and probes targeting different genes to those used in the BioMark^TM^ assay (**Table 2**) (Kim et al. 2008; Rocafort-Ferrer et al. 2022). There are five known genotypes for *T. equi* (designated A-E) circulating in Europe, so we wanted to be sure that we could detect all of those five genotypes if present (Nagore et al. 2004; Bhoora et al. 2009; Salim et al. 2010; Qablan et al. 2013). The Takyon™ No ROX Probe 2X MasterMix Blue dTTP (Eurogentec, Belgium) was used with a final reaction volume of 20 µL containing 10 µL of Master Mix 2X (final concentration 1X), 5 µL of RNase free water, 1 µL of each primer (0.5 µM), probes (0.25 µM), and 2 µL of DNA template. The reaction was carried out using a thermal cycler according to the following cycles: 3 min at 95°C, 45 cycles at 95°C for 10 sec, 55°C for 30 sec and 72°C for 30 sec. Positive controls for both *R. aeschlimannii* and *T. equi* were prepared using a recombinant plasmid from the TA cloning® kit (Invitrogen, USA). A 10-fold serial dilution of the plasmid (from an initial concentration of 0.5x10^8^ copy number/μL) was used to generate standard positive plasmids from 2.5x10^5^ copy number/μL to 2.5x10^-1^ copy number/μL. Samples were detected in duplicates and quantified using the standard plasmids. For *R. aeschlimannii*, we considered negative samples whose Cq number was higher than Cq 37. This detection limit was established regarding the last dilution of the standard curve that could be detected by qPCR. For *T. equi*, most samples were close to or below the detection limit established with the *T. equi* standard curve. Because the protozoan *T. equi* may be circulating at low levels, we included all positive samples.

**Table 2:**
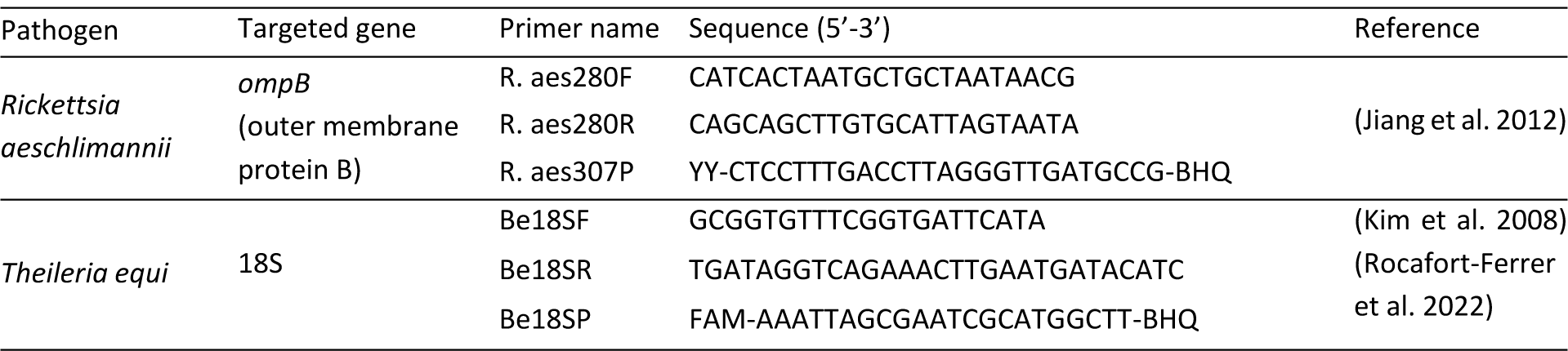
*R. aeschlimannii* and *T. equi* primers and probes sequences used for the detection and quantification method by duplex real-time quantitative PCR.

### Statistical analyses

Statistical analyses were performed with R software 4.2.0. Generalized linear mixed effect models (glmer, package lme4, (Bates et al. 2015)) were used to evaluate the effect of variables (geographic cluster, tick sex, engorgement status, site of collection and host of the tick) on the presence/absence (binomial distribution) of *R. aeschlimannii*, *A. phagocytophilum*, *Francisella*-LE (BioMark^TM^ data) and *T. equi* (qPCR data). In addition, the influence of the geographic cluster, tick sex, engorgement status and the site of collection were also assessed on the loads (qPCR data) of the positive samples for *R. aeschlimannii* and *T. equi* using a glmer with a gamma distribution.

The presences and loads of each pathogen were analysed according to three types of models as follow: A first was used on the whole dataset to assess the influence of the geographic cluster (Aude/Pyrénées-Orientales) and Hérault/Gard), the tick sex (male and female) and the site of collection as a random effect.

A second model was used to evaluate the influence of the engorgement status, with a subset including females only, since the engorgement status of males could not be evaluated. Maximal models included the engorgement status, the geographic cluster the site of collection as a random effect.

Finally, a third model was used to assess the influence of the host on a subset corresponding to ticks located in the Aude/Pyrénées-Orientales cluster, since ticks were collected on both horses and cattle, while they were only collected from on horses in Hérault/Gard. Maximal models included the host, the tick sex and the site of collection as a random effect.

Minimal models and significance of variables were assessed using the ‘ANOVA’ procedure within the package ‘car’ which performs a type III hypothesis (Fox and Weisberg 2018). Post hoc tests were conducted using the function ‘emmeans’ (Tukey HSD test). P-value associated to the random effect “site” was assessed by log-likelihood test in the first model.

Because a very few ticks were positive for *A. marginale*, only descriptive analyses were presented in **Table 3**. We decided to include *Francisella*-LE in the analyses as a control, since this bacterium is definitely identified as a *H. marginatum* primary endosymbiont.

**Table 3:**
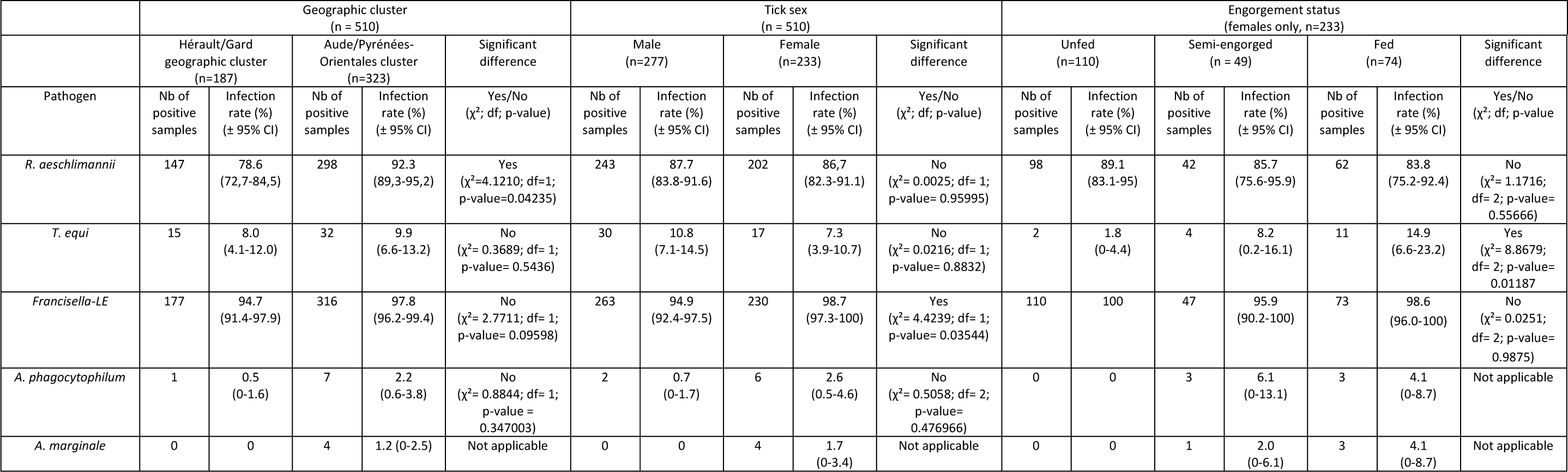
Multivariate analyses of pathogen infection rates. Counts, infection rates and statistical analyses for each variable and each pathogen are presented. For the analyse of the geographic cluster and tick sex, infection rates were calculated by taking into account the number of positive samples for each pathogen and each variable category and the total number of ticks (n=510). The engorgement status influence was analysed on females only (n=233). The number of ticks for each level of variables are indicated for information. « Not applicable » is indicated when no statistical analysis was conducted because of a too low positive sample number. Data were generated using the BioMark^TM^ assay for *R. aeschlimannii*, *Francisella*-LE, *A. phagopcytophilum* and *A. marginale. T. equi* data were generated using the qPCR assay. df is the degree of freedom and X², the Chi-square value.

### Maternal transmission of *R. aeschlimannii*

To assess the potential maternal transmission of the bacteria *R. aeschlimannii*, five fed females collected on the field were individually placed in 50 mL Falcon tubes in an incubator at a temperature of 27°C for several weeks in the BSL3 facility. About 100 eggs belonging to each of five females were isolated and frozen at -80°C for subsequent analyses while the rest of the eggs were left into the incubator. Only a few eggs (from one adult female) hatched into larvae (n=12). The eggs and larvae were placed separately in a 0.2 mL microtube with 100 µL of DMEM and crushed against the bottom of the tube using a sterile needle. DNA was extracted using 100 µL of homogenate with the Genomic DNA tissue kit (Macherey-Nagel, Germany). Detection and quantification of *R. aeschlimannii* was performed by targeting the *ompB* gene by qPCR (**Table 2**).

## RESULTS

### Genotyping of microbes detected in *H. marginatum*

On the 510 ticks analysed using the BioMark^TM^ assay, 445 samples were positive for *R. aeschlimannii*. The sequences obtained were blasted and all allowed the identification of *R. aeschlimannii*; the longest sequence (GenBank Accession Number: PP236764) showed 100% identity with *R. aeschlimannii* collected from a *H. marginatum* from England in 2019 (AN: MT365092.1). The comparison of the obtained sequence with another one from a tick collected on a horse in 2019 in the Gard department of the Occitanie region showed 99% query cover and 98.76% identity (Bernard et al. 2024) (AN: PP379722). The sequencing results allowed the identification of *R. aeschlimannii* and these results were extrapolated to the other samples exhibiting identical amplification patterns.

Two species of *Anaplasma* were detected: eight samples for *A. phagocyotophilum* and four for *A. marginale.* Three sequences from samples positive for *A. phagocytophilum* were blasted and all allowed the identification of *A. phagocytophilum*; the longest sequence (AN: PP265050) showed 99% identity with a sequence isolated from an *Ixodes ricinus* tick in Italy in 2008 (JQ669948.1). Finally, all four sequences of positive samples for *A. marginale* allowed the identification of *A. marginale*, of with the longest sequence (AN: PP218690) showed 100% identity with a sequence obtained from an infected cattle in Iran (GenBank AN : MK016525.1).

*T. equi* was detected in nine samples. One sequence was obtained (AN: PP227163) and blasted resulting in 100% identity with a sequence of *T. equi* from an infected *Rhipicephalus bursa* (MK732476.1) in Corsica, France. *T. orientalis* was detected in one tick (AN: PP358744), the sequence showed 100% identity with a sequence from a *Haemaphysalis longicornis* tick in China in 2014 (MH208633.1).

### Infection rate of *H. marginatum* microbes

Among the 510 ticks analysed with the BioMark^TM^ assay and qPCR, 11.8% [95% CI: 9.0 – 14.6%] were infected with at least one pathogen except *R. aeschlimannii* (*A. phagocytophilum*, *A. marginale, T. equi* or *T. orientalis*). Infection rates using the BioMark^TM^ assay for *A. phagocytophilum* and *A. marginale* were respectively 1.6% [0.5 – 2.7%] and 0.8% [0.02 – 1.6%]. *T. equi* and *T. orientalis* infection rates were respectively 1.8% [0.6 – 2.9%] and 0.6% [0 – 0.6%]. Finally, *R. aeschlimannii* and *Francisella*-LE infection rates reached respectively 87.3% [84.4 – 90.2%] and 96.7% [95.1 – 98.2%].

Re-evaluated *R. aeschlimannii* and *T. equi* infection rates by qPCR were respectively 89.4% [86.7 – 92.1%] and 9.2% [6,7 – 11.7%]. *R. aeschlimannii* infection rate was similar with respect to the detection method used whereas the re-evaluation of *T. equi* infection rate using the qPCR was about five times higher.

Only one co-infection was reported between *A. marginale* and *T. orientalis* in one tick whose infection rate was 0.2% [0 – 0.6%]. *Francisella*-LE was not considered in co-infections neither *R. aeschlimannii* whose pathogenic status will be discussed.

### Multivariate analyses of *H. marginatum* microbes

#### Spatial patterns

##### According to the geographic clusters

The tick-borne pathogen dynamics were firstly assessed by defining a spatial structure linked to the geographic distribution of ticks which consists of two separate areas in the departments Hérault/Gard (HG) and Aude/Pyrénées-Orientales (APO) (**Figure 1**).

1. *R. aeschlimannii* infection rate was significantly influenced by the geographic cluster (χ²= 4.1210; df= 1; p-value= 0.04235, **Figure 2A**), it was higher in the cluster APO: 92.3% [89,3 – 95,2%] compared to the cluster HG: 78.6% [72,7 – 84.5].
2. *T. equi*, *Francisella*-LE and *A. phagocytophilum* infection rates were not significantly influenced by the geographic cluster (**Table 3**).

**Figure 2:**
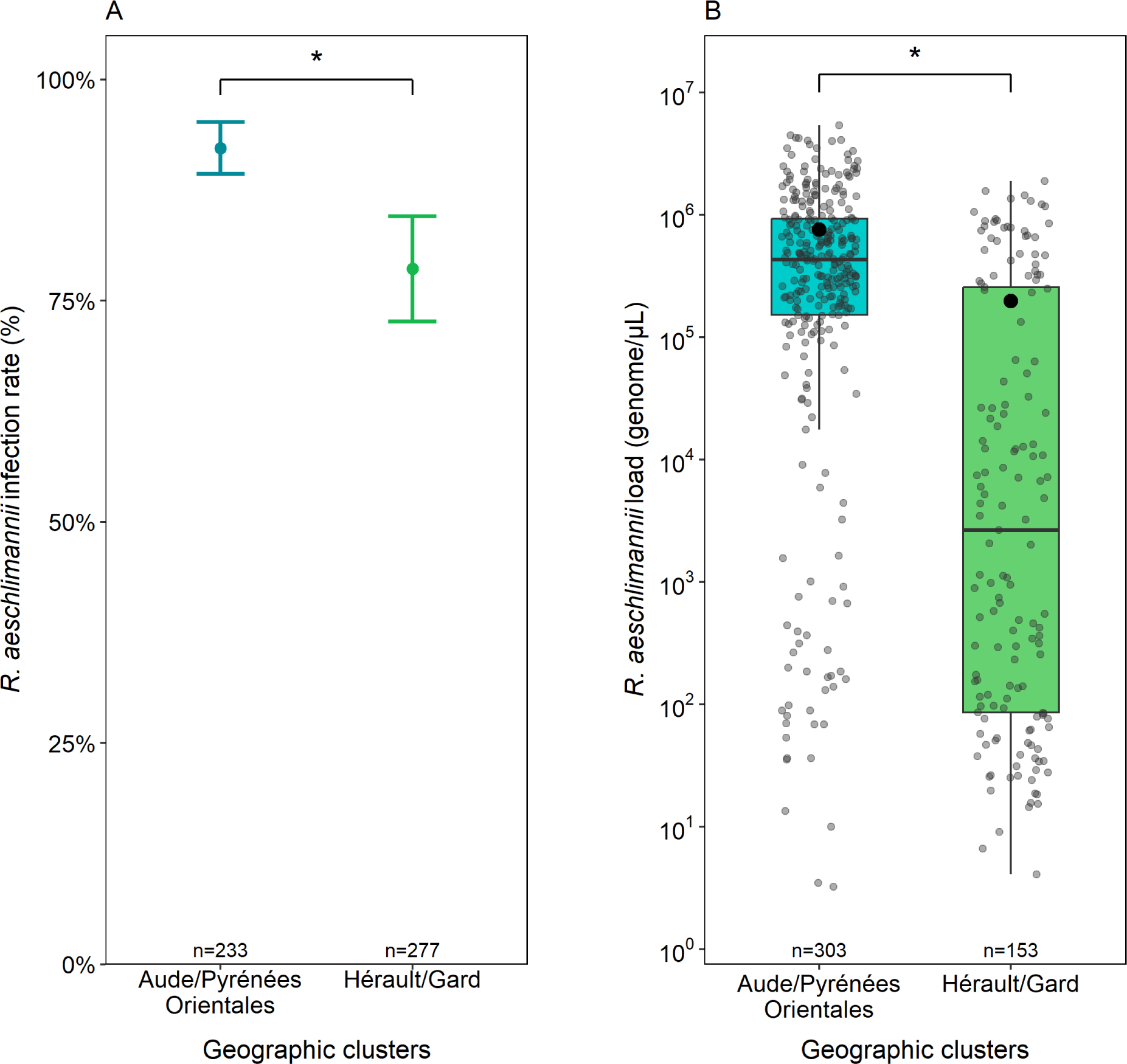
Infection rate and bacterial load of *R. aeschlimannii* in *H. marginatum* ticks collected in the two geographic clusters Hérault/Gard and Aude/Pyrénées-Orientales. The number of ticks per cluster is indicated above the x axis legends. Significant differences are represented by asterisks (*: p-value < 0.05). A. *R. aeschlimannii* presence (1) or absence (0) for each individual tick examined by the BioMark^TM^ assay is summarized by the mean (infection rate in %) and errors bars represent 95% confidence interval. **B.** Bacterial loads expressed in genome.µL^-1^ obtained by qPCR assay are represented by boxplot summarizing the median, 1st and 3rd quartiles. Each grey dot represents the loads of *R. aeschlimannii* for one tick. Mean is symbolized by the black dot.

The influence of the geographic cluster on *R. aeschlimannii* and *T. equi* loads was also analysed.

1. *R. aeschlimannii* bacterial loads were significantly influenced by the geographic cluster (χ²= 6.5292; df= 1; p-value= 0.01061, **Figure 2B**). Indeed, bacterial loads were significantly higher in APO: 7.6x10^5^ genome.µL^-1^ in average [6.5x10^5^ – 8.6x10^5^ genome.µL^-1^] compared to HG: 2.0x10^5^ genome.µL^-1^ in average [1.4x10^5^ – 2.6x10^5^ genome.µL^-1^].

While *T. equi* infection rates were not influenced by the geographic cluster, this spatial variable significantly influenced the *T. equi* loads (χ²= 5.4868; df= 1; p-value= 0.01916) (**Table 4**) as the parasite loads were significantly higher in APO: 1.0x10^3^ genome/µL in average [0 – 3.1x10^3^ genome.µL^-1^] compared to HG: 5.3 genome.µL^-1^ in average [2.9 – 7.7 genome.µL^-1^].

**Table 4:**
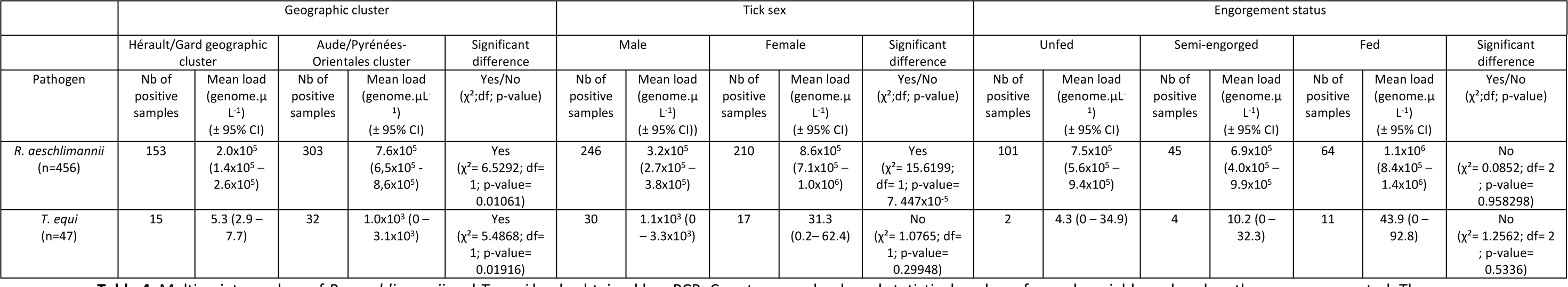
Multivariate analyse of *R. aeschlimannii* and *T. equi* loads obtained by qPCR. Counts, mean loads and statistical analyses for each variable and each pathogen are presented. The mean loads were calculated by taking into account only positive samples for *R. aeschlimannii* (n=456) and *T. equi* (n=47). The engorgement status influence on the pathogens load was analysed on positive females for *R. aeschlimannii* (n=210) and *T. equi* (n=17). The df is the degree of freedom and X², the Chi-square value.

#### According to the collection sites

The collection site had a significant influence on *R. aeschlimannii* infection rate (p-value = 3.88x10^-29^) as illustrated by a variability ranging from 29% to 100% between one site to another in Occitanie. In the cluster HG, 50% of the sites had an infection rate up to 100% compared to 84% of the sites in the cluster APO. Interestingly, infection rates for sites located in the geographical cluster HG were more variable than those from the cluster APO (**Figure 3A, 3B**). The collection site did significantly influence *T. equi* infection rate (p-value= 3.05x10^-8^), ranging from 0% to 41% between one site to another in Occitanie (**Figure S1A**). Finally, it did not have a significant influence on *Francisella*-LE (p-value= 0.707) and A*. phagocytophilum* (p-value= 0.055) infection rates although they varied respectively from 80% to 100% and 3.3% to 20% (**Figure S1B, S1C**). Finally, *A. marginale* infected rates were quite similar between the two sites of collection where it was detected (3.1% and 8.8%) (**Figure S1D**).

**Figure 3:**
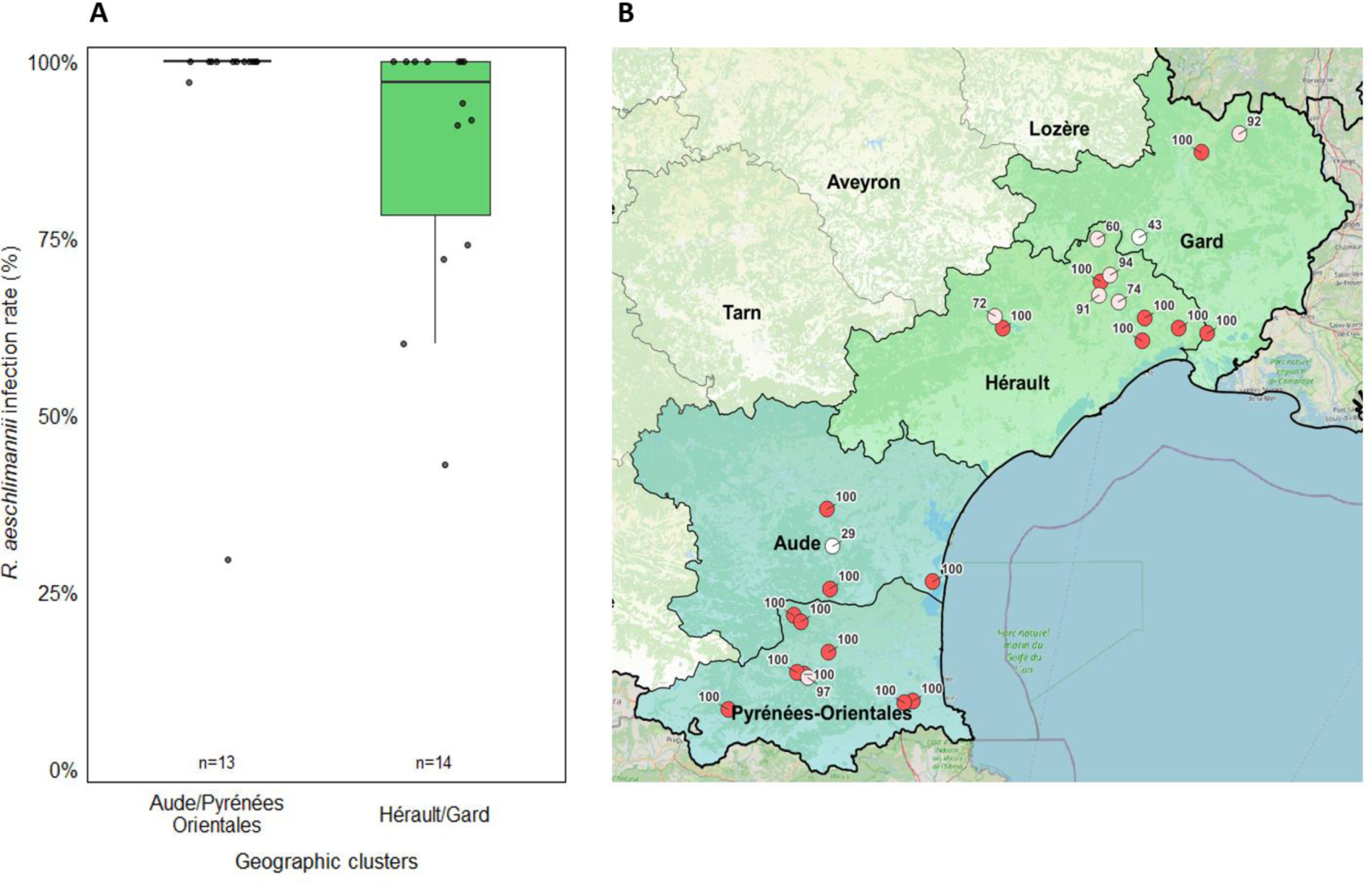
A. *R. aeschlimannii* Infection rate in *H. marginatum* for each site of collection in the two geographic clusters Hérault/Gard and Aude/Pyrénées-Orientales. The number of sites is indicated above the x axis legends. Infection rates for each site were determined with the BioMark^TM^ assay and are represented by black dots. B. *R. aeschlimannii* spatial distribution and its infection rates for each site; represented by a red circle when the infection is 100% and in white when it is less than 100%. Precise infection rates are indicated next to each site.

#### The sex of the ticks

While infection rates estimated for *R. aeschlimannii*, *T. equi* and *A. phagocytophilum* were not significantly influenced by the sex of the ticks (**Table 3**), slight but significant differences were observed for *Francisella*-LE between males and females (χ²= 4.4239; df= 1; p-value= 0.03544) as *Francisella*-LE infection rate was significantly higher in females: 98.7% [97.3 – 100%] compared to males 94.9% (95% CI: 92.4 – 97.5%].

Finally, even though the sex of ticks did not influence the *R. aeschlimannii* infections rates (Table 4, **Figure 4A)**, significant differences were observed for *R. aeschlimannii* loads (χ²= 15.6199; df= 1; p-value= 7.447x10^-5^, **Figure 4B**). Bacterial loads were significantly higher in females: 8.6x10^5^ genome.µL^-1^ in average [7.1x10^5^ – 1.0x10^6^ genome.µL^-1^] compared to males: 3.2x10^5^ genome.µL^-1^ in average (CI: 95% 2.7x10^5^ – 3.8x10^5^ genome.µL^-1^). By contrast, *T. equi* loads were not significantly impacted by the sex of the ticks (**Table 4**).

**Figure 4:**
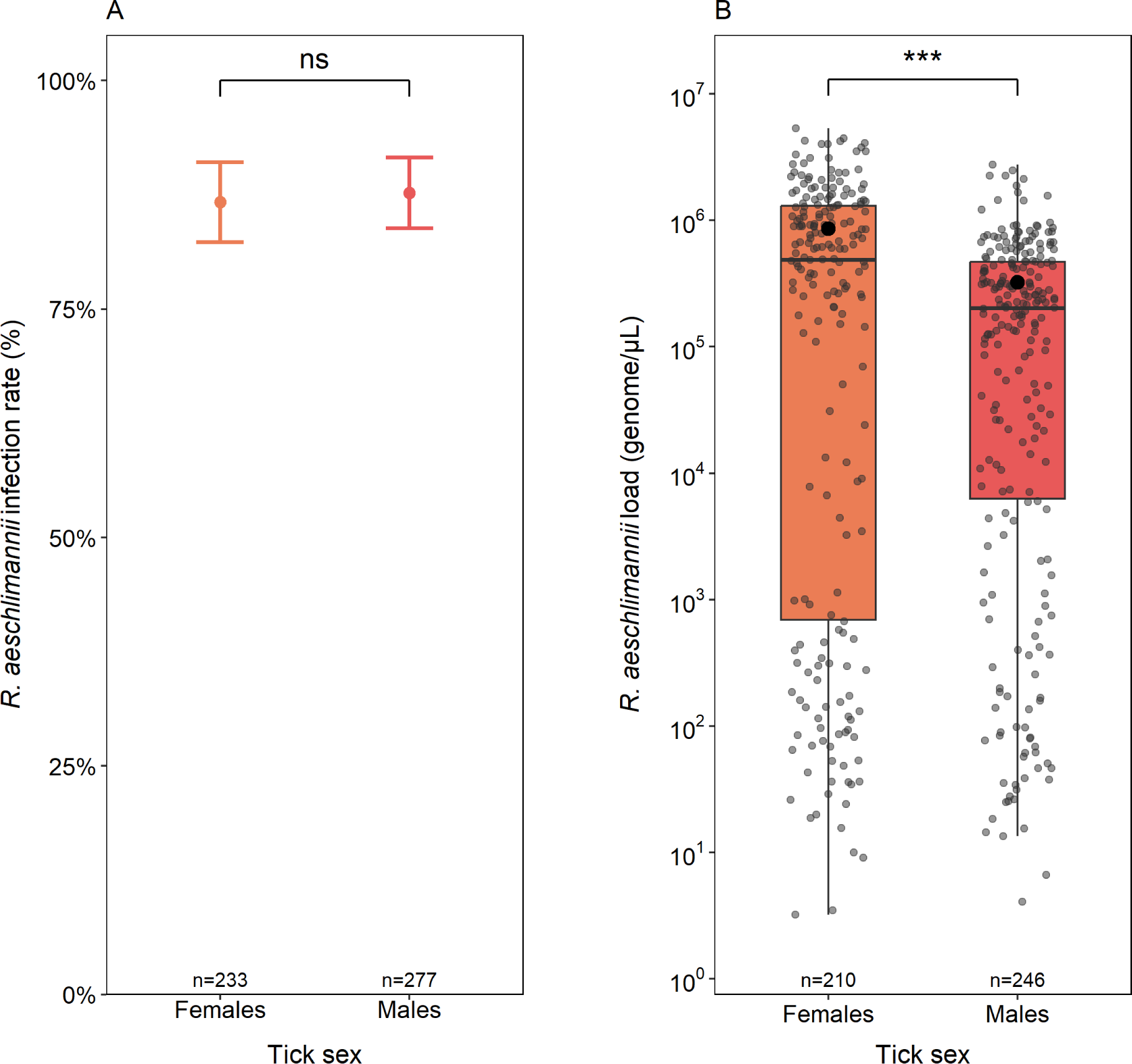
Infection rate and bacterial load of *R. aeschlimannii* in *H. marginatum* males and females. The number of males and females is indicated above the x axis legends. Significant are represented by asterisks (*** : p-value < 0.001). “ns” : non-significant. **A.** *R. aeschlimannii* presence (1) or absence (0) for each individual tick examined by the BioMark^TM^ assay is summarized by the mean (infection rate in %) and errors bars represent 95% confidence interval. **B.** Bacterial loads expressed in genome.µL^-1^ were obtained by qPCR assay and presented by boxplot summarizing the median, 1st and 3rd quartiles. Each grey dot represents the loads of *R. aeschlimannii* for one tick. Mean is symbolized by the black dot.

#### The engorgement status

The infection rate of *T. equi* in female ticks was significantly influenced by the engorgement status (χ²= 8.8679; df= 2; p-value= 0.01187). It was higher in fed females compared to the unfed ones with values reaching 1.8% in average [0 – 4.4%] for unfed females and 14.9% in average [6.6 – 23.2%] for fed females.

Finally, *Francisella*-LE and *R. aeschlimannii* infection rates as well as *R. aeschlimannii* and *T. equi* loads were not significantly influenced by the engorgement status (**Table 3 and 4**). No statistical analysis of the impact of the engorgement status on *A. phagocytophilum* could be performed for due to the a low number of positive samples for this pathogen (**Table 3).**

#### The Hosts

No influence of the tick host was observed on the infection rates of *R. aeschlimannii* (χ²= 0.028; df= 1; p-value= 0.86715), nor on *T. equi* (χ²= 0.0184; df= 1; p-value= 0.892123). The host did show a significant influence on *A. phagocytophilum* infection rate (χ²= 5.6442; df= 1; p-value= 0.01751).

*A. phagocytophilum* infection rate was higher in cattle compared to horses. In APO, 6/7 positive ticks were collected on cattle and 1/7 on horses. No statistical analysis was conducted for *A. marginale* because of a too low positive samples but 4/4 *A. marginale* positive ticks were collected on cattle.

### Maternal transmission of *R. aeschlimannii*

1. *R. aeschlimannii* was detected in all five egg-laying *H. marginatum* females with bacterial loads estimated to 2.0x10^6^ genome.µL^-1^ in average [0 – 5.7x10^6^ genome.µL^-1^]. 100% of the egg pools, belonging to each egg-laying female, were positive for *R. aeschlimannii* with bacterial loads estimated to 3.3x10^5^ genome.µL^-1^ in average [2.2x10^5^ – 4.5x10^5^ genome.µL^-1^]. The only one pool of 12 larvae was also positive for *R. aeschlimannii* with bacterial load estimated to 1.1x10^5^ genome.µL^-1^.

## DISCUSSION

Recently established in the south of France and known to potentially transmit human and animal pathogens, *Hyalomma marginatum* might represent a future problem for both public and animal health. A better assessment of the risk linked to this tick firstly requires an exhaustive identification of *H. marginatum*-borne pathogens and a better characterization of their dynamics. In this context, the objective of this study was to characterize the influence of spatial patterns on both the infection rates and loads of *H. marginatum*-borne pathogens in the region Occitanie in France.

In our study, we detected *R. aeschlimannii*, *T. equi*, *T. orientalis*, *A. phagocytophilum* and *A. marginale* in adult *H. marginatum* ticks. The global infection rates of these pathogens across the Occitanie region were 87.3%, 9.2%, 0.2%, 1.6% and 0.8% respectively. Most of the detected pathogens corresponded to microbes known to have circulated in this geographical area over the last five years (Bernard et al. 2024) which would suggest a certain stability in the circulation of the main *H. marginatum*-borne pathogens in these departments. At first sight, this would suggest identifying sentinel sites with regular monitoring of pathogens transmitted *by H. marginatum*. However, we did not detect the bacteria *Ehrlichia minasensis* previously reported at very low prevalence (in a single tick collected from a horse in the Occitanie region (Bernard et al. 2024). On the other hand, we detected *Theileria orientalis,* responsible for benign theileriosis in cattle (Watts, Playford, and Hickey 2016), in a fed female collected on a bovine in a single site in Pyrénées-Orientales. This detection was particularly unexpected as this parasite was not previously reported in the area. Furthermore, pathogen infection rates were variable across the sites. In epidemiological terms, these results underline the regular monitoring (e.g. multi-year surveys) of *H. marginatum*-borne pathogens at fine spatial scales in order to detect pathogens circulating insidiously in the studied area. Please refer to Bernard et al. 2024 for more details on the vectorial competence discussion of the detected pathogens in *H. marginatum*.

Excluding *R. aeschlimannii* due to the very high infection rate estimated in ticks for this bacterium (see below for specific discussion on *R. aeschlimannii)*, 11.8% of ticks were positive for at least one of the other tested pathogens and only one co-infection was identified between *A. marginale* and *T. orientalis* (0.2%). This low number of co-infection contrasts with *Ixodes ricinus* which has up to five different co-infections (Moutailler et al. 2016). Co-infection are known to enhance disease severity as this is the case for babesiosis and Lyme disease (Grunwaldt, Barbour, and Benach 1983). Overall, while *H. marginatum* is known to carry a wide range of pathogens (Bonnet et al. 2023), we detected a low diversity of pathogens circulating in ticks in the region Occitanie with two *Anaplasma* species, two *Theileria* species, one species of *Rickettsia* and only one co-infection. *H. marginatum* carries fewer pathogens than other tick species of public and veterinary importance like *Ixodes ricinus,* due to differences in their life cycle such as the number and the host spectrum (Lejal et al. 2019; Rizzoli et al. 2014). However, this substantial number of positive ticks for pathogens known to potentially affect both human and animal health, does incites to accentuate the prevention and maintain a regular surveillance of *H. marginatum*-borne pathogens in this area.

### The scheming dynamics of *Rickettsia aeschlimannii*

*Rickettsia aeschlimannii* is known as the agent of spotted fever, a human infection that mainly occurs in North and South Africa, in Greece, Italy and Germany. This bacteria was detected for the first time in *H. marginatum* in 1997 in Morocco (Beati et al. 1997). In our study, *R. aeschlimannii* infection rates were very high whatever the detection method and the targeted gene, the citrate synthase gene through the BioMark^TM^ assay (87.3%) or with the *ompB* gene through qPCR (89.4%). These results are consistent with previous studies performed on *H. marginatum* in southern France which reported high infection rates; from 75% in individual ticks in *H. marginatum* collected around the French Mediterranean sea between 2016 and 2019 (Bernard et al. 2024) to 100% of the pools of *H. marginatum* in Corsica (Grech-Angelini et al. 2020). Such high infection rates observed in our study and many others (Maitre et al. 2023; Bernard et al. 2024) raise the question of the pathogenic status of this detected *Rickettsia,* since a high number of human patients suffering from rickettsiosis should be observed, but reported cases remain extremely rare and only imported cases have been reported in France to date. While Grech-Angelini et al. (2020) suggested that human exposure to *R. aeschlimannii* infection in Corsica is high and human cases of tick-borne spotted fever acquired in Corsica could often be due to *R. aeschlimannii*, there is no information available for its vectorial competence based on experimental approaches, showing that this assumption must be nuanced. Further studies are required to characterize the full genome of *R. aeschlimannii* found in *H. marginatum* ticks in order to compare it with the complete genome of *R. aeschlimannii* isolated from a human case. This will help to determine whether the bacterium encountered in ticks is the one responsible for human rickettsiosis cases or whether it is a different strain, potentially involved in a symbiotic relationship with the tick.

Several of our results raise the question of whether *R. aeschlimannii* is a symbiotic bacterium in *H. marginatum*. Indeed, the transovarian transmission of *R. aeschlimannii* in *H. marginatum* paired with its high infection rates both support this assumption (Parola, Paddock, and Raoult 2005; Azagi et al. 2017). In the literature, such hypothesis have already been suggested for *H. marginatum* (Maitre et al. 2023; Boularias et al. 2021). Tick symbionts are typically classified as primary or secondary symbionts, depending on their role in the tick’s survival. In the present case, *R. aeschlimannii* infection rate is high (87.3%), but still not reaching typical infection rate of primary symbiont close to 100% like this is the case for *Francisella*-LE (96.7%). In addition, its status as a primary symbiont seems unlikely because of the significant spatial variability in terms of infection rates and loads, where the infection rate of *Francisella-*like endosymbiont is very stable regardless of spatial patterns. Therefore, *R. aeschlimannii* would rather be a secondary endosymbiont. Overall, this hypothesis contrasts with the postulate that *Rickettsia* symbionts are most common in ticks of the genera of *Ixodes*, *Amblyomma*, and *Dermacentor*, whereas it has been less frequently found in *Rhipicephalus*, *Haemaphysalis*, and *Hyalomma* ticks (Burgdorfer 1981; Hussain et al. 2022; Nováková et al. 2018).

Regarding the functions of symbiotic *Rickettsia* in tick physiology, it was already demonstrated that they can be involved in the nutrition, such as the *Rickettsia buchneri* endosymbiont in *Ixodes scapularis* or *I. pacificus* (Tokarz et al. 2019; Benson et al. 2004), as it harbours all required genes for folate biosynthesis (Hunter et al. 2015). This nutritive role is unlikely in the case of *H. marginatum,* insofar as *R. aeschlimannii* do not have complete biosynthesis pathways for B vitamins, as opposed to the two primary symbionts *Francisella*-LE and Midichloria that possess either complete biotin, riboflavin or folic acid pathways (Buysse et al. 2021).

Other function of *Rickettsia* in arthropods in the literature mention defence (Łukasik et al. 2013) and reproductivity (Engelstädter and Hurst 2009). Our results indicate similar infection rates whatever the tick sex assuming that the function of *R. aeschlimannii* appears to benefit both male and female ticks (whether fed or not). Its role in reproductivity seems unlikely since higher proportion should be observed in females. In regard of the present information, we suggest that *R. aeschlimannii* would be involved in tick defence against abiotic factors and further investigations for example using transcriptomics approaches should be conducted in the future to confirm such a hypothesis.

In the present study, strong spatial patterns were observed for *R. aeschlimannii*, whether it be at a large spatial scale (geographic cluster) or at a small one (from one site to another, especially in Hérault/Gard). Some of the infection rates variability between the sites might be explained by unequal tick numbers per sites. It is also possible that while *R. aeschlimannii* is present in ticks since it is maternally transmitted, the quantities are too low to be detected efficiently, contributing to this variability. In this study, we thus propose two hypotheses to explain this uneven spatial distribution of *R. aeschlimannii* infection rates.

The first is to assume that *R. aeschlimannii* infection rate can be influenced by environmental characteristics (humidity, temperature, vegetation, host variability). *Rickettsia aeschlimannii* would replicate preferentially when environmental conditions threaten the tick and this would be in Aude/Pyrénées-Orientales rather than in Hérault/Gard. In the literature, it was reported that infection by *Rickettsia* sp. of the spotted fever group in *Ixodes ricinus* is influenced by the geographic location and their environmental characteristics (forest fragmentation, vegetation, hosts) (Halos et al. 2010; Narasimhan et al. 2021). Another study reported significant difference in loads of *Rickettsia* phylotype G021 between *Ixodes pacificus* from different collection sites and vegetation habitats (Cheng et al. 2013). In order to explore this hypothesis, it would be interesting to evaluate the infection rate by *R. aeschlimannii* and the loads from ticks experimentally maintained in microcosms located in different environmental conditions (vegetation, temperature) in the field (Rahajarison et al. 2014), in order to evaluate the influence of these variables.

The second hypothesis is that the presence and loads of *R. aeschlimannii* would vary from one *H. marginatum* population to another. It is interesting to note that, the spatial distribution of *R. aeschlimannii* in the two geographical areas (Hérault/ Gard and Aude/Pyrénées-Orientales) is consistent with two genetically differentiated populations of *H. marginatum* characterised using mitochondrial markers clusters main haplotypes (Giupponi et al., personal communication). The presence of structured genetic differentiation could be explained by the introduction of tick populations by migratory birds or horses and cattle from different geographical origins. These introduced tick populations could harbour microbial communities with different compositions, which could explain the spatial patterns of *R. aeschlimannii* in French ticks.

### Dynamics of other *H. marginatum* microbes

As expected, *Francisella*-LE was unsurprisingly detected in almost all ticks (96.7%). *Francisella*-LE is a known primary symbiont vital for tick survival and reproduction by assuring the production of B vitamins production (Duron et al. 2018; Azagi et al. 2017). Our results are consistent with previous studies in which infection rates by *Francisella*-LE reached 97% in *H. marginatum* ticks collected in southern France (Bernard et al. 2024) and 90% of *H. marginatum* pools collected in Corsica (Grech-Angelini et al. 2020). The absence of influence of spatial groups on the rate of infection is not surprising, as it is a primary symbiont necessary for the tick’s physiology. On contrast, the sex of ticks had a slight but significant influence on the presence of *Francisella*-LE with higher infection rates in females than in males. It is interesting to note that this observation has already been reported in *Dermacentor* ticks (Sperling et al. 2020; Dergousoff and Chilton 2012). Although *Francisella*-LE quantification was not estimated in our study, this difference may also be explained by the fact that *Francisella*-LE loads might be lower in males and therefore make the detection more difficult, as already demonstrated in *D. variabilis* ticks (Chicana et al. 2019). This might be linked to the functional role of this symbiont in the reproduction, which is probably less necessary for males than for females (Duron et al. 2017). Neither the engorgement status of females nor the host of the tick had any influence on the presence of *Francisella*-LE.

Unlike *Francisella*-LE*, A. phagocytophilum* was detected in only 1.6% of the collected ticks.

*A. phagocytophilum* is the etiological agent of human granulocytic anaplasmosis (HGA), a tick-borne fever in ruminants and equine granulocytic anaplasmosis (Chen et al. 1994; Karshima et al. 2023; Stannard et al. 1969). *Anaplasma phagocytophilum* is maintained in wild ruminants (roe-deer, white- tails deer, white-footed mice), domestic animals (cattle, sheep) and birds in an enzootic cycle which is consistent with our results since 7/8 of *A. phagocytophilum* infected ticks were collected from cattle. Since *A. phagocytophilum* is considered an emerging pathogen of horses (Dzięgiel et al. 2013), it is not surprising to find a tick collected from a horse infected by this bacterium. Finally, most of the infected ticks were females (6/8) that were either semi-engorged or fed although no significant influence of the tick sex or the engorgement status was observed. This result would indicate that ticks become infected after a blood meal suggesting that the tick infection is a reflect of the host infection. However, such a hypothesis might imply that more ticks should be infected. One thing is for sure that the role of *H. marginatum* as a vector of this pathogen is questioned and further studies need to address its vectorial competence (Bernard et al. 2024)

Interestingly, another *Anaplasma* species*, Anaplasma marginale,* was detected in ticks with an infection rate of 0.8%. *Anaplasma marginale* is responsible for fever, anaemia, weight loss and abortions in cattle. The main reported vectors of this bacteria are *Rhipicephalus* spp. and *Dermacentor* spp. in tropical and subtropical regions that can be found in Europe. Statistics were not applicable due to the low number of *A. marginale* infected ticks (n=4) but descriptive data showed that all infected ticks were engorged females (semi or fully fed). All the ticks were collected in two sites of Aude/Pyrénées-Orientales, on cattle. The most likely hypothesis is that the four female fed ticks infected with *A. marginale* became infected by blood feeding on the same infected animal bovine host.

Finally, the equine piroplasmosis agent *T. equi* parasite was detected in our study. We first estimated *T. equi* infection rate at 1.8% with the BioMark^TM^ assay. However, this approach made us aware of the probable underestimation of the infection rate as we used primers that did not allow the detection of all genotypes potentially circulating in France. We re-evaluated *T. equi* infection rates using another gene (18S) that can target several genotypes of *T. equi* by qPCR an infection resulting in a rate of 9.2%. Interestingly, this infection rate was very low compare to a study that showed 43% of *H. marginatum* infected with *T. equi* collected from horses in Camargue, next to the Occitanie region (Rocafort-Ferrer et al. 2022). This large difference can probably be explained by the design of the tick collection that aimed to target stables where cases of piroplasmosis are frequently reported. Most of *T. equi* infected ticks were collected on horses (33/47). As horses are known to be reservoirs of *T. equi* and a high circulation rate of piroplasmosis has been described in France, it is not surprising that the majority of infected ticks were collected on horses (Nadal, Bonnet, and Marsot 2022). However, it should be noted that 14/47 of the infected ticks were collected from cattle, which is more surprising since cattle are not known to be susceptible to infection by *T. equi* but rather to *T. annulata* and

*T. parva*. Such a result could suggest maternal transmission of this parasite in ticks but this has not yet demonstrated and should be tested experimentally in *H. marginatum.* It could also suggest that the adult tick could have been infected during its immature stages on a susceptible host. However, it is currently unknown if the hosts of immature stages (birds and lagomorphs) can carry and be infected with *T. equi*.

While *T. equi* infection rates were not influenced by the geographic cluster, significant differences were observed on *T. equi* loads, with a lower number of genomes.µL^-1^ in Hérault/Gard than in Aude/Pyrénées-Orientales. However, this result has to be nuanced regarding the fact that a tick collected on cattle in a site of Aude/Pyrénées-Orientales presented very high *T. equi* loads compared to other infected ticks.

Finally, although *T. equi* infection rates and loads were not influenced by the sex of the tick, *T. equi* infection rate was significantly higher in fed females compared to unfed ones, suggesting that *H. marginatum* ticks tend to be infected by *T. equi via* the blood meal on a parasitemic horse rather than being responsible for the transmission of *T. equi* to an uninfected horse.

## CONCLUSION

This study characterized the spatial distribution of *H. marginatum*-borne pathogens in the region Occitanie where the tick has recently become established. At the scale of the whole region, we reported that (i) 11.8% of *H. marginatum* ticks were infected with at least one species of *Theileria* or *Anaplasma*; (ii) the infection rate of *R. aeschlimannii* was very high (87.3%), questioning its pathogenic status regarding the low number of human cases; (iii) *R. aeschlimannii* is hypothesized to be a secondary symbiont of *H. marginatum* due to its high infection rates and its maternal transmission; (iv) infection rates of all detected pathogens were quite variable from one site of collection to another, demonstrating the importance of the sampling effort for pathogen circulation surveillance. The particularly marked spatial patterns for *R. aeschlimannii* presence and loads might be linked to the tick population. Beyond the spatial scale, pathogens dynamic can also fluctuate depending on the temporal scale, even at a monthly scale (Pollet et al. 2020; Lejal et al. 2019). It is necessary to highlight temporal patterns that might affect pathogens and symbionts in *H. marginatum*. To do this annual and monthly monitoring of ticks collected in a given location would help us identify temporal dynamics.

## Acknowledgments

The authors are grateful to Maud Marsot, Facundo Munoz, Maxime Prat and Haoues Alout for their helpful advice on statistics. We are grateful to Mickaël Mège and Claire Bonsergent for technical assistance and to Frédéric Stachurski for the relevant discussions.

The work was funded by the Holistique project (défi clé RIVOC Occitanie region, University of Montpellier): “*Hyalomma marginatum* in Occitanie region: analysis of biological invasion and associated risks”.

## Author contribution

**Conceptualization:** TP, SM, CJK. **Funding acquisition**: TP. **Methodology**: CJK, CG, DB, CR, SM. **Formal analysis**: CJK. **Field and resources**: CJK, CB, HJP, KH, IK, LM, FS, LV, TP. **Writing-original draft**: CJK, TP, SM. All authors read and approved the final manuscript.

## Supplementary material

**Figure S1:**
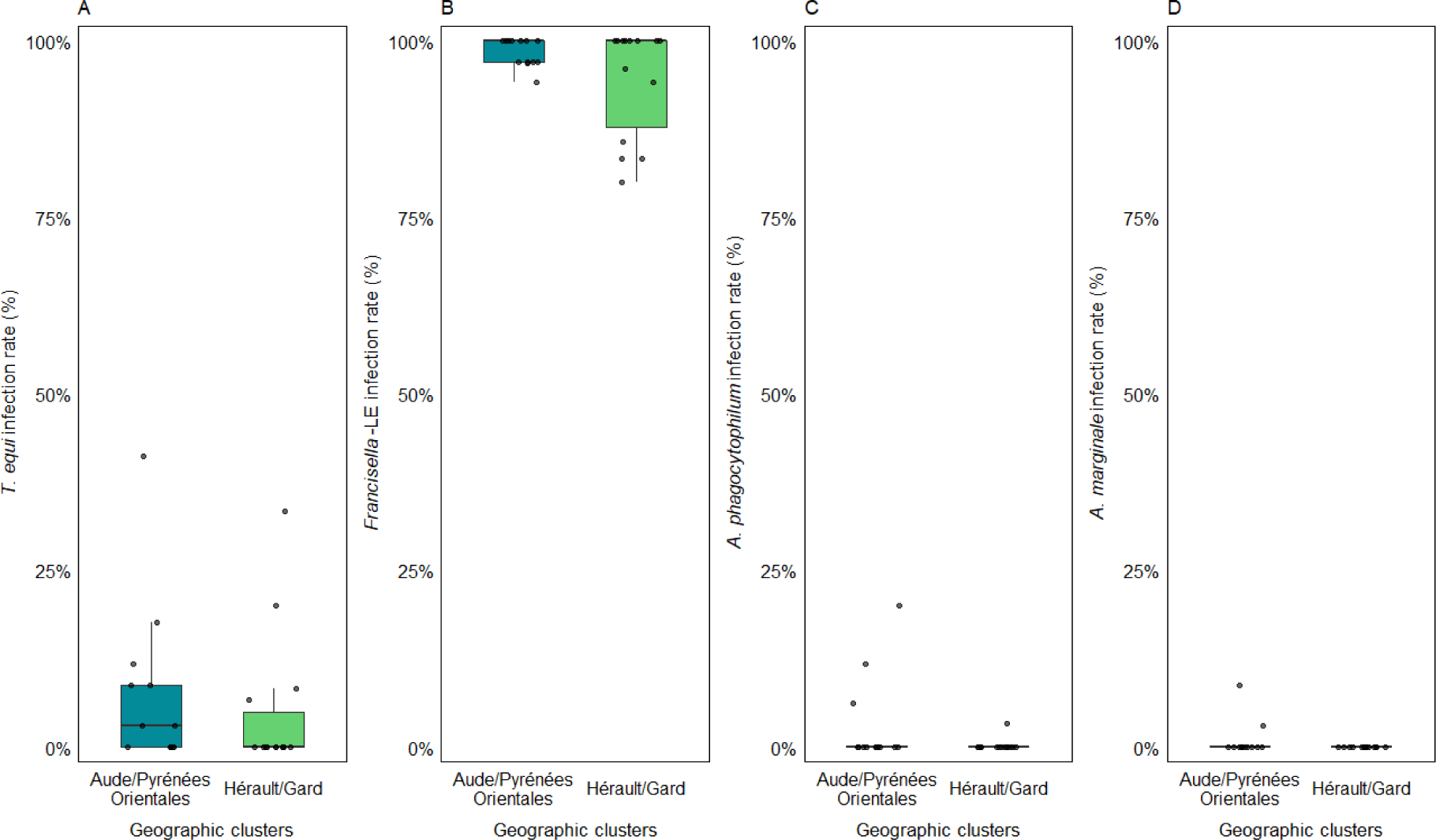
Infection rates of *T. equi* (A), *Francisella*-LE (B) *A. phagocytophilum* (C) and *A. marginale* (D) in *H. marginatum* ticks for each site of collection in the two geographic clusters Hérault/Gard (n=14 sites) and Aude/Pyrénées-Orientales (n=13 sites). Infection rates for each site represented by black dots were determined with the BioMark^TM^ assay for *Francisella*- LE, *A. phagocytophilum* and *A. marginale* and with the qPCR assay for *T. equi*.

**Table S1:**
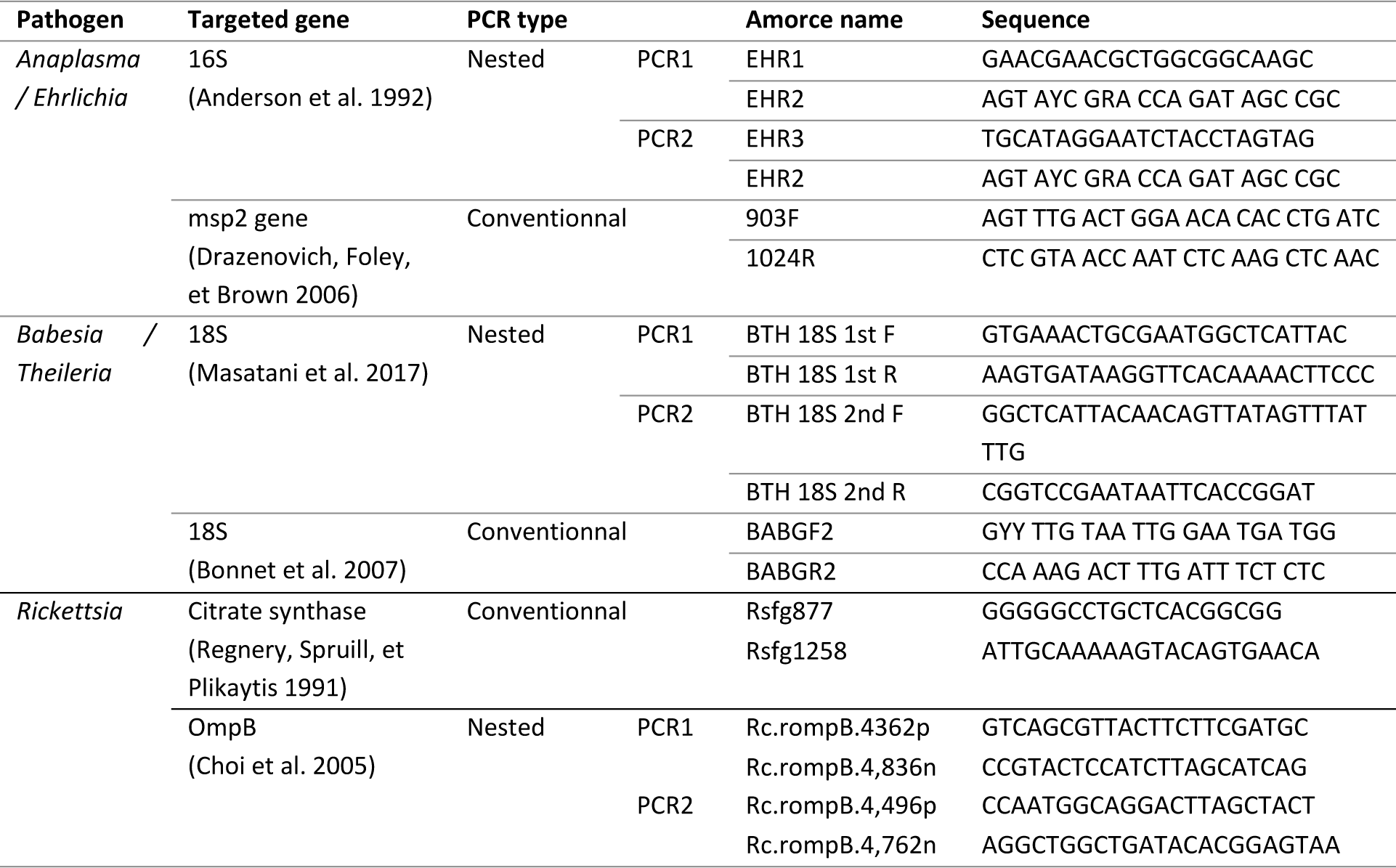
Tick-borne pathogens primers and probes sequences used for detection confirmation by PCRs and qCPRs.

